# Optimizing a human fecal assay that elicits bacterial swarming

**DOI:** 10.1101/794487

**Authors:** Arjun Byju, Deeti Patel, Weijie Chen, Sridhar Mani

**Author notes:** Correspondence: Sridhar Mani, MD (Co-correspondence: Arjun Byju), Albert Einstein College of Medicine, 1300 Morris Park Avenue, Chanin Building, 302-D1, Bronx, NY, USA, 10461, OR. Equal Contribution (Arjun Byju, Deeti Patel).

## Abstract

A distinct property of many bacteria is swarming: swift movement across a surface through flagella propulsion. Early research indicates that bacterial swarming can be a protective host response to intestinal inflammation. Central to the further study of bacterial swarming in human health is an effective and replicable assay for swarming that can accommodate complex material, such as fecal matter. To date, nearly all swarming assays described in the literature are specific for bacteria grown in culture, most often *Pseudomonas*. In this paper, we describe a protocol for discerning swarming of bacteria from frozen human fecal samples. Moreover, we tested 4 variables that may influence the effectiveness of the assay: the method by which frozen samples were thawed, the concentration of agar used in the Lysogenic broth (LB) agar plate, the volume of LB agar poured in the plate, and the volume of sample inoculated. We found that while the type of thaw and volume of LB agar had little to no effect on swarming, greater concentrations of agar were negatively correlated with swarming and greater volumes of the sample were positively correlated with swarming.

## Introduction

A distinct property of many bacteria is swarming—swift movement across a surface through flagella propulsion [1, 2]. While this behavior can provide certain bacteria with a competitive advantage in their specific ecosystem [3] the larger impact of swarming on host organisms remains unclear. Recently, it has been posited that swarming may be a protective host response to intestinal inflammation; early research indicates that bacterial swarming is a specific biomarker of intestinal distress and may be used clinically to diagnose and predict flare-ups of Inflammatory Bowel Disease, Crohn’s Colitis, and Ulcerative Colitis, among others [see manuscript under submission, Chen et. al].

Nevertheless, the ultimate clinical application of such findings relies on the existence of a consistent and reproducible assay for bacterial swarming that accommodates complex human material, such as fecal matter. While various groups have created assays of swarming motility, these protocols are largely restricted to the study of single strains of bacteria and in culture, namely *Pseudomonas* [4-7].

This paper addresses this lacuna by describing a protocol for assaying swarming from frozen human fecal samples. Specifically, covariates are identified as being integral to the consistency of the assay, taking as a starting point variable that putatively influence the occurrence of swarming in cultured *Pseudomonas* (thickness of plate, percentage of agar) [5,7] as well as volume of sample inoculated and method of thaw.

## Methods and Materials

Human fecal samples were obtained in 2015 (under protocols IRB# 2009-446 and 2015-4465) and since have been preserved in −80°C freezers. One sample, p-12, came from a patient without intestinal distress, and as a known non-swarmer was inoculated on each plate as a negative control. The other sample, p-17, came from a patient with significant intestinal inflammation, and was our experimental group. Our “standard protocol” consisted of preparing a 0.5% Agar Lysogenic Broth (100 mL H_2_O, 1 g Tryptone, 0.5 g Yeast Extract, 0.5 g NaCl, 0.5 g Agar). After autoclaving all ingredients, plates were poured to a volume of 20 mL, and at least one hour later were inoculated with sample. Samples were removed from the freezer and thawed. Then, we inoculated p-12 onto one half of the plate and p-17 onto the other, dried it for 10 minutes under biological hood, and scanned an image of the plates. This image is the “before image.” Then, plates were placed upright in an incubator, set to 37 °C with a humidity of 80% RH. After 24 hours plates were removed, assessed in a binary fashion for the presence or absence of swarming, and scanned for an “after picture.” Swarming was defined visually by observation of a leading swarming edge and a surfactant ring around the edge of the colony. The folds of expansion were tabulated from these before and after pictures using the NIH’s ImageJ program (version 1.59e) and were calculated as [(final area - initial area)/initial area]. From these values we also calculated a “swarm score” for each colony of 1, 2, or 3. Scores of 1 include expansion folds of 0-5, scores of 2 include expansion folds of 5-25, and scores of 3 include expansion folds greater than 25.

Using this “standard protocol” we focused on evaluating 4 major variables: (a) type of thaw from frozen, (b) volume of sample inoculated, (c) percentage of agar in plate, and (d) total volume of LB agar poured in the plate. (a) The type of thaw was differentiated by either thawing the samples on ice for 1 hour (n = 35) or thawing them on the bench, at ambient temperature, for 8 minutes (n = 38). In both cases, samples were completely liquid upon completion of thaw. (b) We tested inoculation volumes of 2.5 μL (n = 20), 5.0 μL (n =10), and 7.5 μL (n = 20). (c) We measured different agar concentrations all following the above “standard protocol” merely with altered levels of agar. For the 0.4% agar plates, 0.4 grams of agar were used in the agar Lysogenic Broth (LB) (n = 10). The 0.5% agar plates contained 0.5 grams of agar (n = 20), the 0.6% agar plates contained 0.6 grams of agar (n = 10) and the 0.7% agar plates contained 0.7 grams of agar (n = 20). All other ingredients remained identical to the standard preparation. (d)We tested swarming on plates with LB agar volumes of 15 mL (n=16), 25 mL (n=16), and 30 mL (n = 16) as a proxy for thickness of plate.

## Results

Agar concentration was found to be inversely correlated with the propensity to swarm. As the agar concentration in the plate increased, the expansion fold of the colonies decreased, showing significant differenence between the lower agar concentrations (0.4% and 0.5%) and the higher concentrations (0.6 and 0.7%) [Figure 1A]. Moreover, the magnitude of swarming, as measured by distribution of the Swarm Score, shows swarming covering a higher area at concentrations of 0.4% and 0.5% than at 0.6% or 0.7% [Figure 1B]. The volume of fecal sample inoculated on the plate was positively associated with swarming. As the volume of the inoculated spot increased from 2.5 to 5.0 to 7.5 μL, the size of the expansion fold over the 24-hour period increased significantly [Figure 1C]. Additionally, greater proportions of level 3 scorers on the Swarm Score were found in the 5.0 μL group and even greater proportions in the 7.5 μL group, indicating that larger amounts of fecal matter leads to swarming that covers a greater area [Figure 1D].

**Figure 1.**
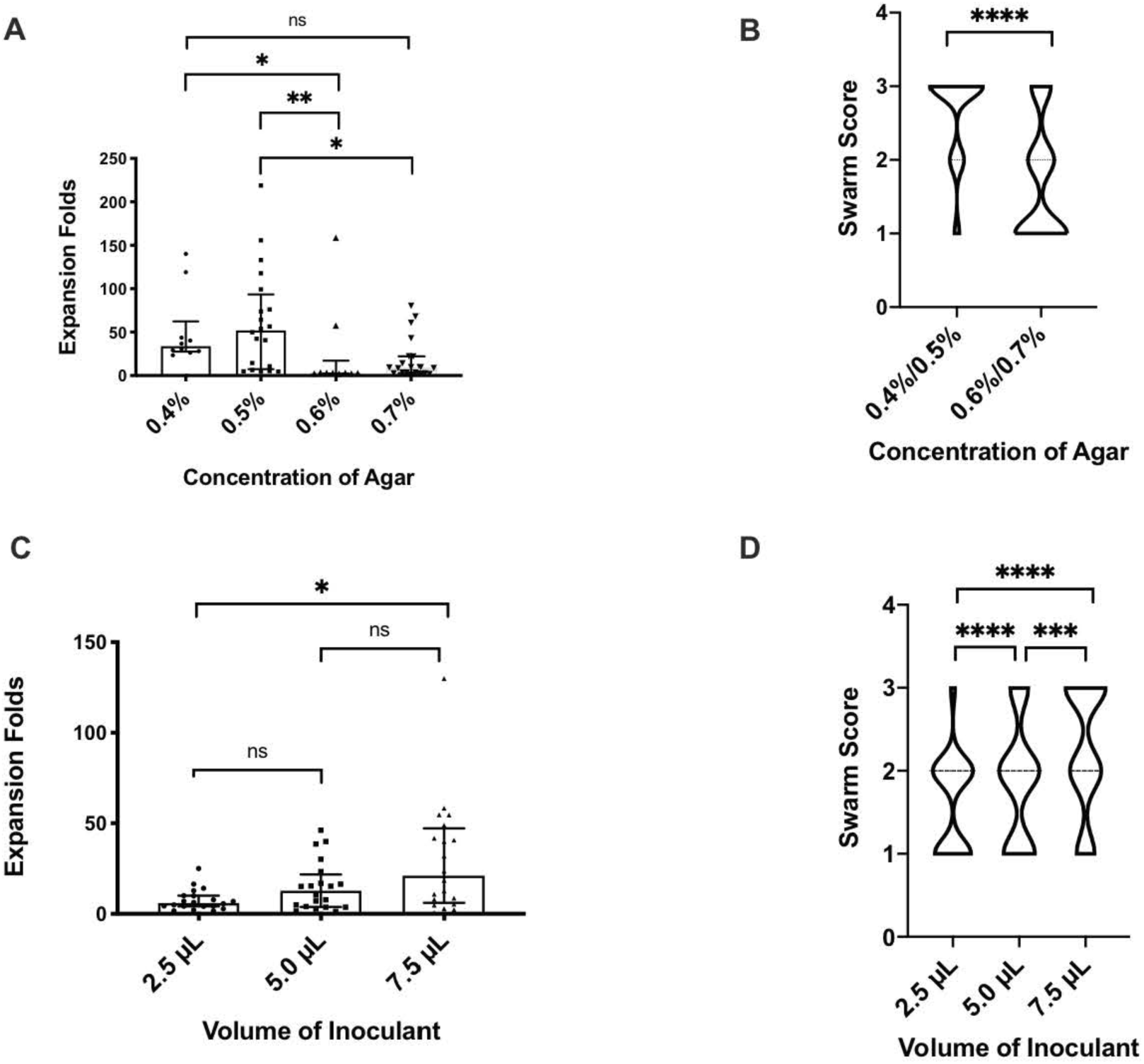
The effects of agar concentration and inoculant volume on fecal sample swarming expansion rate. LB Agar was prepared with varying percentages of agar: 0.4%, 0.5%, 0.6%, and 0.7%. 5 μL of sample were inoculated on 20 mL petri dishes (n = 10, in singlet and duplicate). Simultaneously, 20 mL LB Agar plates were poured and spotted with varying volumes of sample: 2.5 μL, 5.0 μL, and 7.5 μL (n = 10, duplicate). **A.** indicates the expansion folds of the swarm colony in varied agar concentrations, compared to the before picture at t = 0. **B.** indicates the swarm score given to each swarmer with a value of 1, 2, or 3. Scores of 1 include expansion folds of 0-5; scores of 2 include expansion folds of 5-25; scores of 3 include expansion folds greater than 25. A Mann-Whitney test was conducted to show the statistical significance of the inverse correlation between the increased concentration of agar and decreased expansion folds. **C.** indicates the expansion fold of the swarm colony in varied volumes of inoculation, compared to the before picture at t = 0. Data are represented as median and interquartile range, significance tested using the Kruskal-Wallis ANOVA test. Significance results are indicated on the graph. **D.** indicates the swarm score given to each swarmer with a value of 1, 2, or 3. A one sample t and Wilcoxon test were conducted to show the statistical significance of the direct correlation between increased volume of inoculation and increased expansion folds. *\*p < 0.05*; *\*\* p < 0.01; *** p < 0.001; ****p* < *0.0001.* ns, not significant. LB, Lysogenic Broth.

The method of thawing the samples from frozen was found to have a non-significant impact (defined as a response that is less than half-log when comparing methods) on swarming. When samples were removed from the freezer and thawed at room temperature or on ice, the expansion folds did not differ in any significant way, nor did the distribution of the Swarm Score [Figure 2a-b].

**Figure 2.**
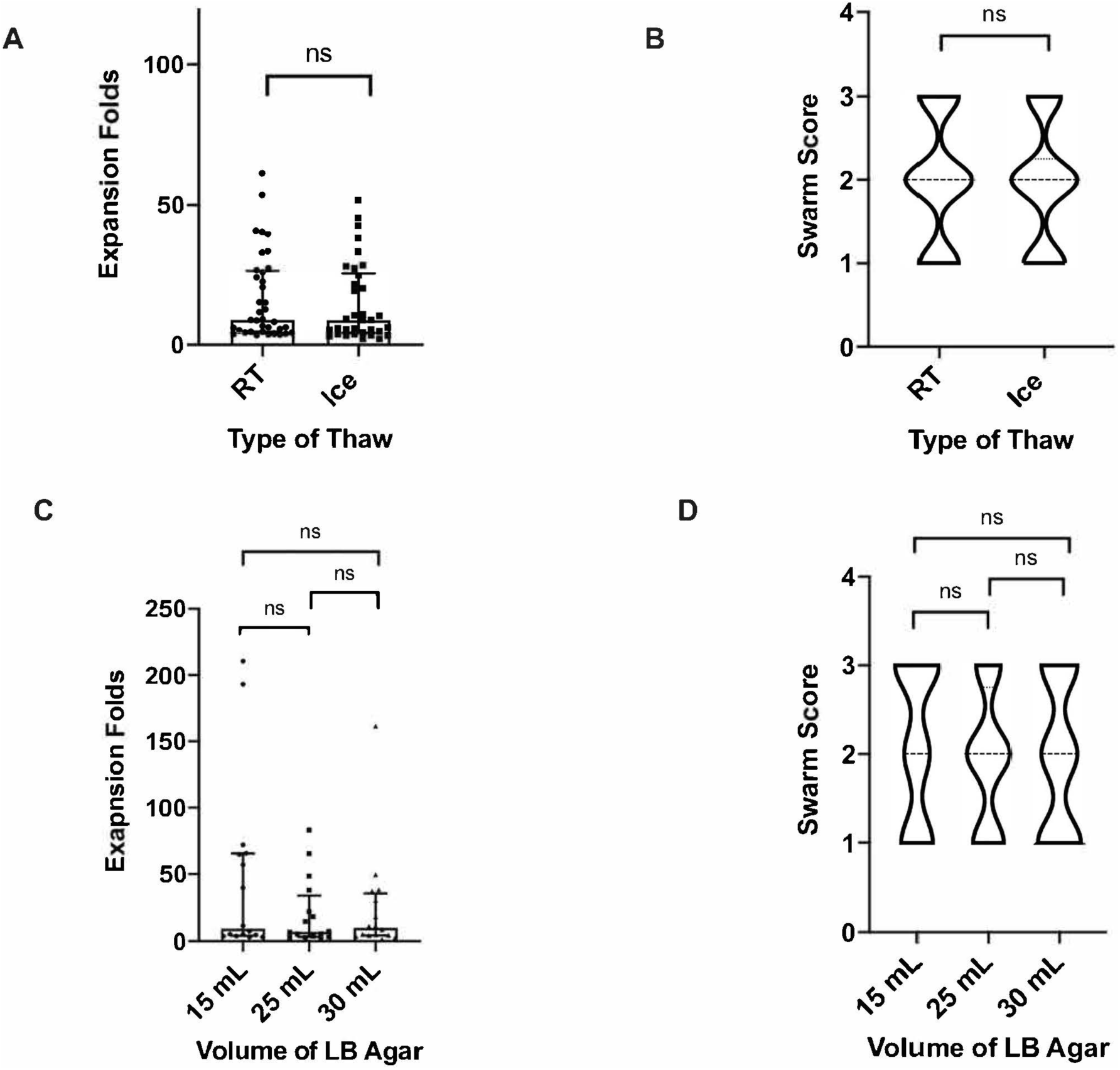
The effect of type of thaw and volume of LB agar on fecal sample swarming expansion rate. Fecal samples were thawed at room temperature or on ice before inoculating on LB agar (n = 10, at least in triplicate). Simultaneously, LB Agar was poured with volumes 15 mL, 25 mL, and 30 mL and 5 μL of sample were inoculated (n = 8, duplicate). **A.** indicates the fold of expansion of the swarm colony after room temperature thaw and after ice thaw, compared to the before picture at t = 0. Data are represented as median and interquartile range, significance tested using Mann-Whitney test. **B.** indicates the swarm score given to each swarmer with a value of 1, 2, or 3. Scores of 1 include expansion folds of 0-5. Scores of 2 include expansion folds of 5-25. Scores of 3 include expansion folds greater than 25. A normality and lognormality test show no significant difference between the type of thaw and the occurrence of each swarm score. **C.** indicates the expansion folds of the swarm colony in varied volumes of LB agar on plate, compared with before picture at t = 0. Data are represented as median and interquartile range, significance tested using the Kruskai-Wallis ANOVA test. **D.** indicates the swarm score given to each swarmer with a value of 1,2, or 3. A Mann-Whitney test was conducted to show that the difference in expansion fold presented between each volume of LB Agar shows no statistical significance, ns, not significant. LB, Lysogenic Broth. RT, room temperature.

Finally, when the volume of the LB agar plate was altered from 15 mL to 25 mL to 30 mL, the fold expansions did not differ significantly, nor did the distribution of the Swarm Score [Figure 2c-d].

## Conclusions

Our findings both concur with and contradict the suggestions for assay optimization posited by those who have studied swarming in cultured bacteria, implying that covariates for the optimal assay of swarming in complex human material, like stool, are unique. First, our finding that agar concentration was inversely linked to swarming aligns with what Tremblay and others [4-7] have identified. It appears that increasing amounts of agar slow the rate of swarming; however, it appeared that colonies on higher agar concentration plates could swarm eventually if given more time. Nevertheless, for researchers with time constraints, using 0.4% agar plates should lead to the fastest results with the minimum usage of reagents, as we saw a 100% identification of known swarmers at this concentration. We would caution against lowering the agar concentration any further as it may allow even characteristically non-swarmers to exhibit plate-covering morphology, making the distinction between swarmers and non-swarmers difficult— and functionally damaging the sensitivity and specificity of the assay.

Secondly, our finding that greater volumes of sample inoculated on the plate produced greater percentages of swarming was predictable, given the nature of the complex material used for sample. With 7.5 μL of sample there is 3 times as much material from which bacteria may be collected to swarm. However, 2.5 μL inoculants tend to also produce swarming if the length of time for observation of swarming is extended. For researchers considering replicating this assay, this presents a trade-off between sparing sample and sparing time. However, most samples captured from human feces should provide more than 7.5 μL, even after multiple aliquots have been made. Furthermore, swarming was more pronounced, due to larger fold expansions, at higher volumes of inoculant—rendering visual identification of swarming easier. Therefore, we suggest using the higher volume of inoculant to ascertain knowledge of swarming within 24 h.

Third, the fact that the type of thaw did not play a significant role in swarming is relevant for clinical application, as we can assume that most fecal samples will be frozen prior to transport from one location to another. At the outset, we had speculated that the modality of thaw may play a role in the presentation of swarming. However, multiple replicates showed no major difference in swarming. The relative inconsequentiality of the type of thaw allows researchers and diagnosticians to save precious time in their ascertainment of swarming from human stool.

Finally, we did not find a marked difference in the ability to observe swarming based on the ‘thickness’ of the plate, contraindicating the findings of Ha et al (5). If greater volume (or thickness) of a plate were to influence swarming, one might expect 15 mL LB agar plates to have the lowest percentage of swarmers and for the swarming percentage to increase steadily with increasing volume. However, this was not the case, with both 15 mL and 30 mL plates producing swarming at rates not significantly divergent from those presented on 25 mL plates. Given this information we again posit that for researchers desiring to use lab resources judiciously, a standard plate volume of 20 mL LB agar should suffice for this assay.

While other covariates may in the future be considered in order to further enhance the effectiveness of this assay, as of now, we believe the protocol to be simple, fast, and reproducible. By using a standard protocol for LB-agar plates and considering either 0.4 – 0.5% agar and/or 7.5 μL of sample, researchers can visualize swarming from frozen, complex human material swiftly and with little difficulty.

## Acknowledgements

Research was conducted under a Student Research Fellowship Award from the Crohn’s and Colitis Foundation of America (#640806), awarded to A.B. The data acquisition was also partially supported by a Grant# 362520 Broad Medical Research Program (BMRP, not Litwin) at CCFA (Crohn’s & Colitis Foundation of America) (to S.M). The authors would like to thank Arpan De, and Hao Li for their advice, suggestions for variables to explore, and help with formatting and editing.

